# Genetic determinants of penicillin tolerance in *Vibrio cholerae*

**DOI:** 10.1101/337949

**Authors:** Anna I. Weaver, Shannon G. Murphy, Benjamin Umans, Srikar Tallavajhala, Ikenna Onyekwere, Stephen Wittels, Jung-Ho Shin, Michael VanNieuwenhze, Matthew K. Waldor, Tobias Dörr

## Abstract

Many bacteria are resistant to killing (“tolerant”) by typically bactericidal antibiotics due to their ability to counteract drug-induced cell damage. *Vibrio cholerae*, the cholera agent, displays an unusually high tolerance to diverse inhibitors of cell wall synthesis. Exposure to these agents, which in other bacteria leads to lysis and death, results in a breakdown of the cell wall and subsequent sphere formation in *V. cholerae*. Spheres readily recover to rod-shaped cells upon antibiotic removal, but the mechanisms mediating the recovery process are not well-characterized. Here, we found that the mechanisms of recovery are dependent on environmental conditions. Interestingly, on agarose pads, spheres undergo characteristic stages during the restoration of rod shape. Drug inhibition and microscopy experiments suggest that class A Penicillin Binding Proteins (aPBPs) play a more active role than the Rod system, especially early in sphere recovery. TnSeq analyses revealed that LPS and cell wall biogenesis genes as well as the sigma E cell envelope stress response were particularly critical for recovery. LPS core and O-antigen appear to be more critical for sphere formation/integrity and viability than Lipid A modifications. Overall, our findings demonstrate that the outer membrane is a key contributor to beta lactam tolerance and suggest a role for aPBPs in cell wall biogenesis in the absence of rod-shape cues. Factors required for post-antibiotic recovery could serve as targets for antibiotic adjuvants that enhance the efficacy of antibiotics that inhibit cell wall biogenesis.

## Introduction

The emergence of antibiotic resistance in bacterial pathogens requires the development of new drugs and novel strategies to combat infection. However, antibiotic resistance is not the sole explanation for antibiotic treatment failures. Instead, some infections are caused by fully susceptible pathogens that are thought to survive antibiotic treatment due to a high level of drug tolerance, i.e., the capacity to stay alive in the presence of otherwise bactericidal drugs (1–4). Dormant persister cells, which resist killing by all available antibiotics (4), represent an extreme form of antibiotic tolerance. However, susceptible (non-persister) bacteria are sometimes capable of surviving severe antibiotic-imposed damage, potentially providing an opportunity to acquire or evolve resistance mechanisms. In addition, surviving bacteria typically exhibit a prolonged lag phase after drug exposure (the post-antibiotic effect), during which they can repair antibiotic induced damage. Currently our knowledge of the molecular processes underlying antibiotic tolerance and the post-antibiotic effect is limited. Understanding the mechanistic underpinnings of post-antibiotic recovery could yield insights enabling the development of novel approaches to target tolerant organisms.

We and others have previously shown that some Gram-negative bacteria, including *Burkholderia pseudomallei, Pseudomonas aeruginosa* and *Vibrio cholerae*, the causative agent of cholera, exhibit high tolerance to ordinarily bactericidal cell wall acting antibiotics (e.g. beta lactams) (5–7). In *V. cholerae*, for example, exposure to beta lactam antibiotics at multiples of the minimum inhibitory concentration (MIC) results in cell wall loss, similar to well-studied model organisms, such as *E. coli* (5). However, in contrast to *E. coli, V. cholerae* survives as wall-deficient spheres, similar to L-forms (8), with the notable difference that *V. cholerae* spheres do not divide while in this wall-less state. Remarkably, however, the wall-deficient spherical cells remain viable and have minimal plating defects on media lacking antibiotics (5). Sphere survival *in vivo* (in the mouse intestine) and *in vitro* is enabled by the two-component cell wall stress response system *wigKR* (aka *vxrAB)*, which controls several processes including cell wall (peptidoglycan, PG) biosynthesis, motility, type VI secretion and biofilm formation (9–11).

The many steps in PG synthesis include cytoplasmic production of the lipid II precursor, translocation of this precursor into the periplasm, and finally precursor incorporation into the cell wall sacculus via polymerization (transglycosylation, TG) and intercrosslinking (transpeptidation, TP) reactions. TG and TP reactions are mediated by two spatiotemporally distinct entities, the Rod system (with RodA as the polymerase and a class B Penicillin Binding Protein [bPBP] as the crosslinking enzyme) and the class A PBPs (aPBPs) that can catalyze both TG and TP reactions (12). In addition, aPBPs require outer-membrane localized lipoproteins (Lpos) for their activity (13, 14) and the Rod system is associated with (and requires for its activity) the cytoskeletal actin homolog MreB (15–18). Almost the entire *V. cholerae* PG synthesis pathway is upregulated through *wigKR/vxrAB* in response to antibiotics that disrupt cell wall synthesis (9), with the notable exception of components of the Rod system.

We have little knowledge of how *V. cholerae* cell envelope biogenesis pathways enable recovery from the antibiotic-induced spherical state and if additional factors contribute to survival and recovery from this state. Here, we have characterized the post-antibiotic recovery process in *V. cholerae*. Microscopy using fluorescent protein fusions and cell wall stains revealed that during an ordered sphere recovery process, aPBP1a localizes prominently to the outgrowth area and its function appears to account for the majority of the initial deposition of new cell wall material. In contrast, the Rod system, which is ultimately required for sphere recovery, plays a minor role in the initial recovery stages. We also used transposon insertion sequencing (TnSeq) to identify the genetic requirements for *V. cholerae* tolerance to penicillin and found that there is an enrichment in genes important for cell wall and outer membrane biogenesis functions among mutations that confer post-antibiotic fitness defects. Collectively, our findings reveal the pleiotropic nature of beta lactam tolerance, provide potential targets for beta lactam adjuvants, and have implications for the role of aPBPs in *de novo* PG template generation.

## Results

### Distinct mechanisms of recovery in different growth conditions

In previous work, we used microscopy to characterize *V. cholerae* sphere formation following exposure to antibiotics that interfere with cell wall synthesis (5). Here we used a similar approach to investigate how spheres revert to rod shape. As observed previously, *V. cholerae* cells grown in minimal medium exposed to penicillin G form non-dividing spheres exhibiting well-defined demarcations between the phase-dark cytoplasm, an enlarged periplasmic space visible as a phase-light bubble, and a clearly visible outer membrane (**Fig. 1A**). Timelapse light microscopy was used to monitor cell morphology on agarose pads after removal of the antibiotic by washing. In these conditions, approximately 10 to 50% of cells fully recovered to form microcolonies (see **Movie S1** for an example). While these conditions were not as favorable for recovery as plating on LB agar (5), they enabled us to discern steps in sphere recovery, which appeared to take place in partially overlapping stages in wild type (wt) cells (**Fig. 1B**). Initially, phase dark material engulfed the periplasmic space (engulfment stage); then, the now elliptical-shaped cells reduced their widths (constriction phase) followed by elongation (elongation phase); finally these elongated cell masses gave rise to rod-shaped cells, which proliferated into a microcolony.

**Figure 1.**
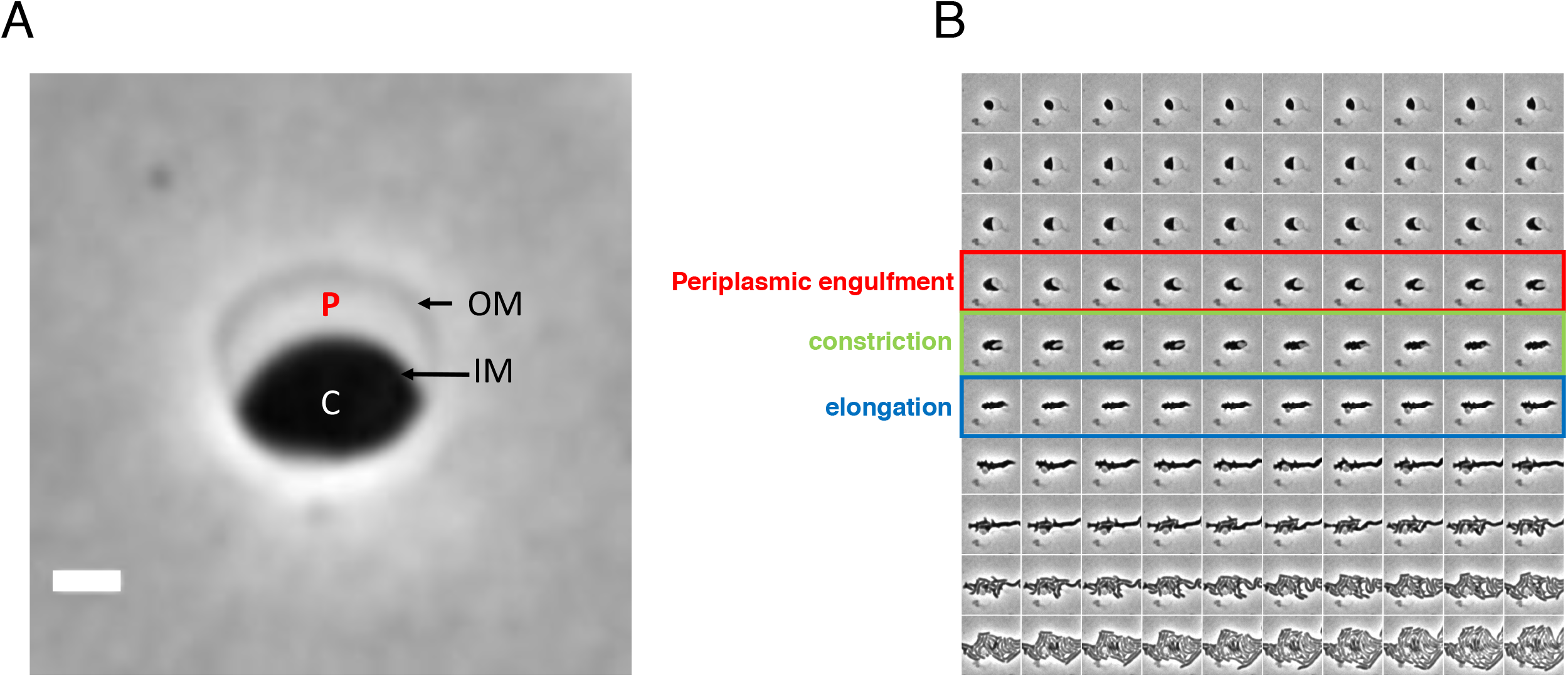
Recovery of *V. cholerae* rod morphology on agarose pads. A) Sphere anatomy after 3 h of treatment with PenG. OM, outer membrane, IM, inner membrane, C, cytoplasm, P, periplasm. Cellular compartments as determined in (5) using fluorescent protein fusions with known localization patterns. Scale bar = 1 μm B) Representative timelapse images of PenG-generated spheres after removal of the antibiotic on an agarose pad.

The pattern of recovery of rod shape described above is distinct from that described for osmostabilized, beta lactam treated *E. coli* cells (19); however the latter experiments were conducted in microfluidic chambers rather than agarose pads. Unlike *E. coli, V. cholerae* does not require osmostabilization for sphere formation; furthermore, *V. cholerae* spheres retain viability and structural integrity in LB and minimal medium, as well as in rabbit cecal fluid (5). Unlike the conditions in microfluidic chambers, agarose pads may provide external structural support to recovering spheres. Consistent with this idea, we found that the pattern and dynamics of recovery were very different when we repeated recovery experiments in liquid M9 minimal medium. Following exposure to penG and washing, cells were intermittently removed from the liquid medium and imaged. We did not observe the distinct stages of recovery observed on agarose pads; in general, sphere morphology did not change for the duration of the experiment (12 h), except for a slight increase in volume (**Fig. 2**). However, normal, rod-shaped cells appeared after ~ 4-5 hours of post-antibiotic incubation (**Fig. 2**, yellow arrow). We surveyed ~ 100 cells per time point in each of two biological replicate experiments and did not find any intermediates, suggesting that if such intermediates form, they do so at a frequency <1/100. The origin of the rod-shaped cells is not clear, but they may have directly budded off of spheres from a newly-formed pole juxtaposed to the periplasm, similar to the recovery protrusions observed in *E. coli* after treatment with beta lactams (19) or lysozyme (20). Indeed, we observed some rods that appeared to be budding off of spheres (**Fig. 2**, red arrow). Thus, the morphological transitions and dynamics of sphere to rod conversion are dependent on specific culture conditions and may rely on distinct mechanisms.

**Figure 2.**
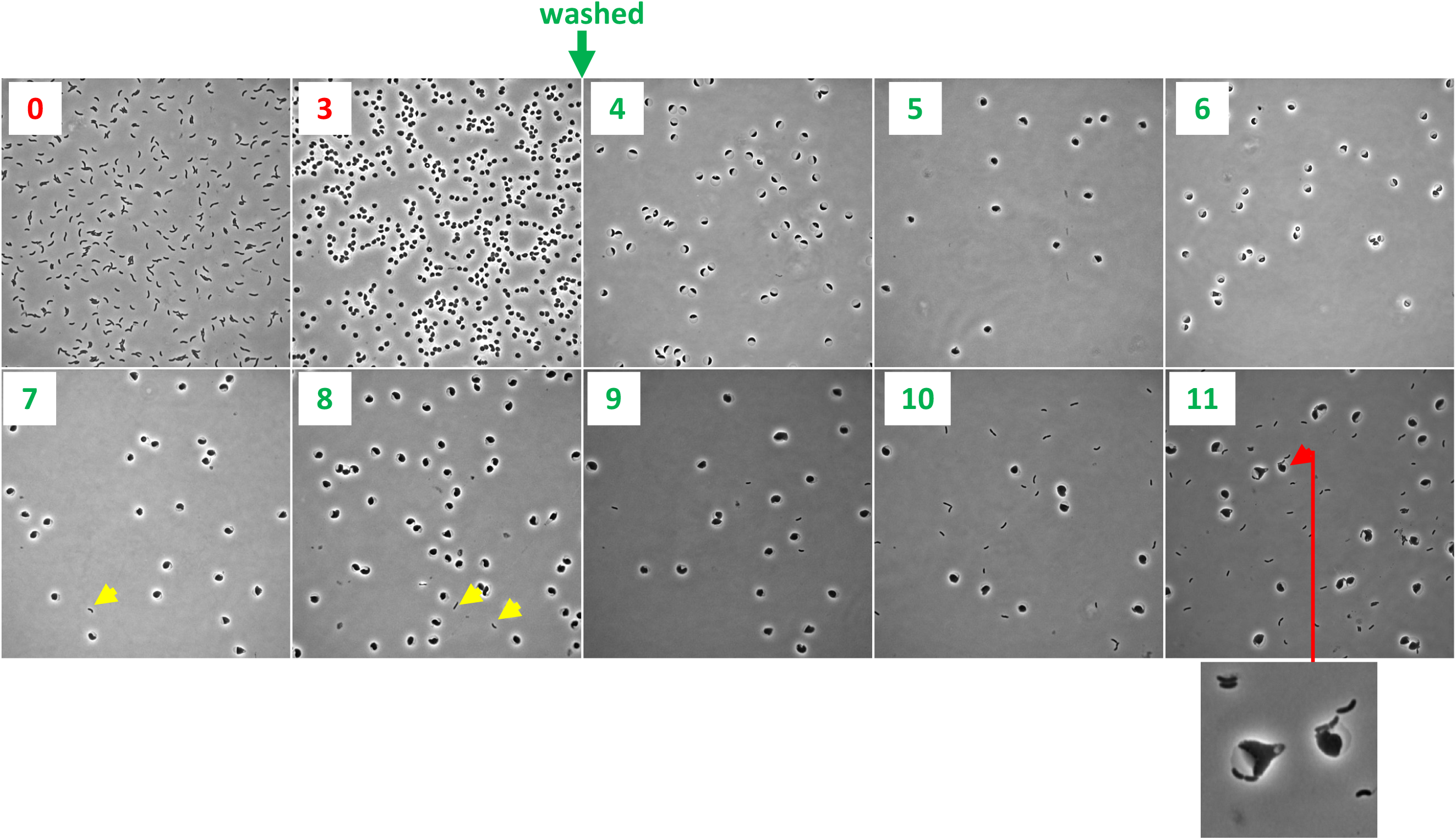
Sphere recovery in liquid medium. Cells were grown to a density of ~ 2 × 10^8^ cfu/ml (T0) in minimal medium, exposed to penicillin G (100 μg ml^−1^, 10x MIC) for 3 h (T3), then washed twice to remove the antibiotic and then imaged every hour. Yellow arrowheads show rod shaped cells and red arrowhead (plus enlarged window) shows sphere apparently budding of a rod.

### PBP localization dynamics during sphere recovery

*V. cholerae*, like the model rod-shaped organisms *E. coli* and *B. subtilis*, encodes two distinct cell wall synthesis machineries, namely the RodAZ-MreBC-PBP2 elongation complex (Rod system) and the aPBPs (15, 21, 22). Nearly the entire cell wall synthesis pathway, including aPBPs, is upregulated by the *wigKR* cell wall stress response two component system. Members of the Rod system, however, are conspicuously absent from the *wigKR* regulon (9). We thus hypothesized that aPBPs were crucial determinants of post-antibiotic recovery.

To investigate the role of PBP1a, *V. cholerae*’s primary aPBP (23) in the recovery process, we created a functional (**Fig. S1**) PBP1amCherry translational fluorescent protein and tracked its localization in recovering spheres on agarose pads. In the first stages of recovery, PBP1amCherry was diffuse, but then it assumed a striking, band-like pattern along the leading edge of the periplasmic engulfment, migrating ahead of the phase-dark cytoplasmic material (**Fig. 3, yellow arrow**). Inhibiting PBP1a’s TG activity using moenomycin (10 μg ml^−1^, 10x MIC), arrested sphere recovery in the pre-engulfment stage and prevented proper PBP1a localization, suggesting that the recovery process depends on PBP1a’s PG synthesis capabilities (or at least transglycosylation function). We also tested whether MreB was necessary for PBP1a’s leading edge localization by treating the recovering PBP1amcherry strain with the MreB inhibitor MP265 (24) (200 μM, 10 × MIC). Inhibition of MreB suppressed recovery and completion of periplasmic engulfment, establishing that MreB is important for sphere to rod recovery as shown before for *E. coli* (19). However, engulfment was only partially defective in spheres treated with MP265 and PBP1a still localized in a concentrated, band-like pattern in the presence of MP265 (**Fig. 3, green arrow**). Thus, while both MreB and PBP1a are important for recovery, aPBPs seems to function earlier than the Rod system in the process.

**Figure 3.**
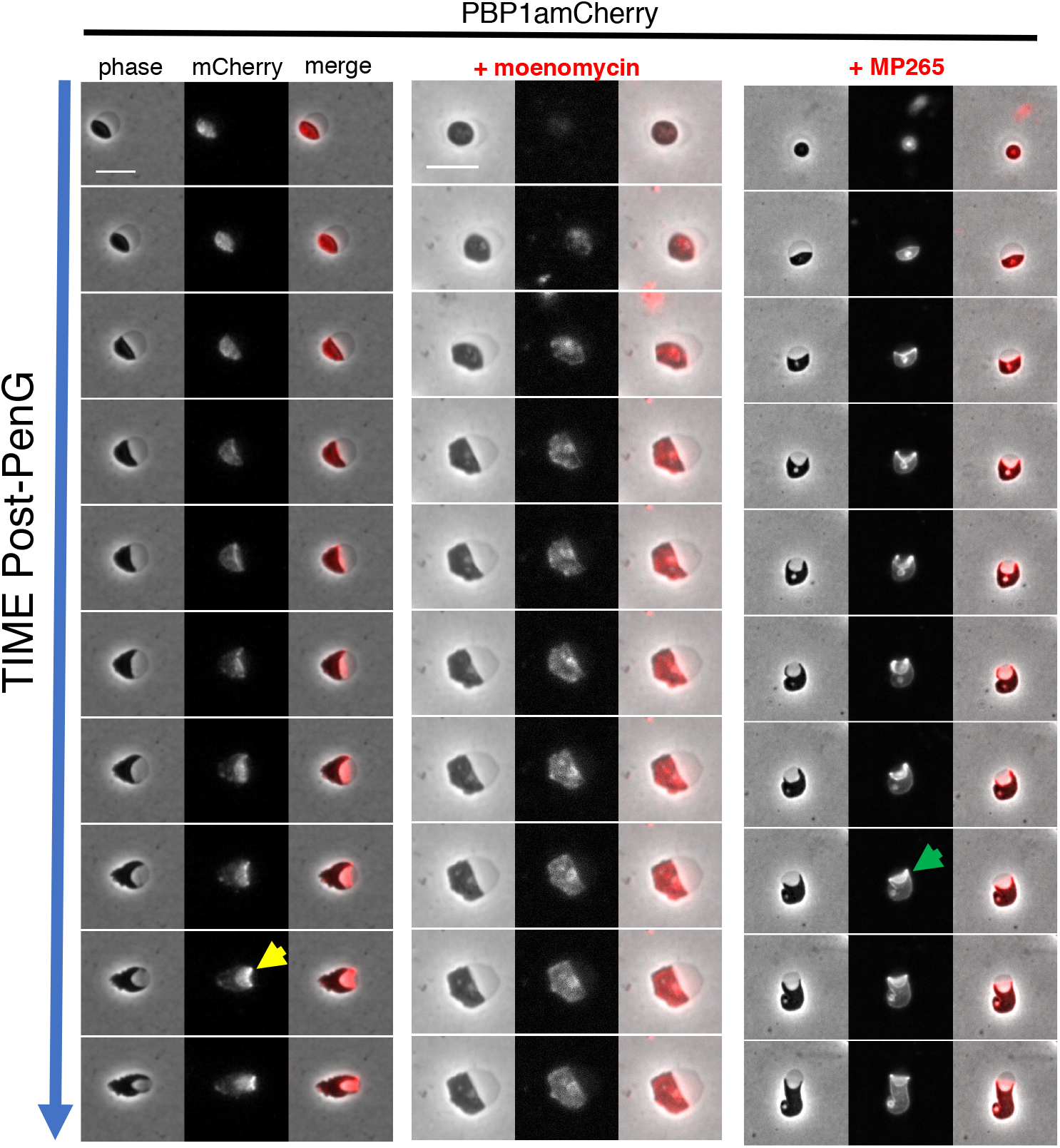
PBP1a localizes to the leading edge of engulfment during sphere recovery. Cells were exposed to PenG (μg ml^−1^, 10 × MIC) for 3 h, followed by washing and application to agarose pads containing either no antibiotic, the aPBP inhibitor moenomcyin or the MreB inhibitor MP265 (both at 10x MIC). Frames are 10 min apart, scale bar = 5 μm. Arrows point to examples of ring-like localization of PBP1a in untreated spheres (yellow arrow) or those exposed to MP265 (green arrow).

### The aPBPs are important for the initiation of cell wall synthesis early in sphere recovery

Since PBP1a was concentrated around the leading edge during the engulfment process, we hypothesized that the aPBPs might be required for the commencement of cell wall synthesis after antibiotic-induced murein degradation. To test this, we treated cells with PenG, removed the antibiotic by washing, and then used the fluorescent D-amino acid HADA to visualize insertion of new cell wall material. An Δ*ldtA* Δ*ldtB* mutant defective in L,D transpeptidase activity (25) was used in these experiments to exclude PG synthesis-independent HADA incorporation. In untreated spheres, cell wall deposition generally started at the opposite side of the periplasm (**Fig. 4**). This is likely the place where aPBPs and their OM activators interact first, as the inner and outer membranes are in close proximity in this area. In the presence of the MreB inhibitor MP265 (at 10 × MIC), initial cell wall deposition was reduced compared to untreated spheres, but remained detectable. In contrast, when cells were incubated with moenomycin (10 × MIC), incorporation of HADA-labeled material was drastically reduced (**Fig. 4**, see **Fig. S2** for image adjusted to lower dynamic range). It follows that while both the aPBPs and MreB are required for sphere recovery, the aPBPs are more active than the Rod system in producing nascent PG in recovering spheres.

**Figure 4.**
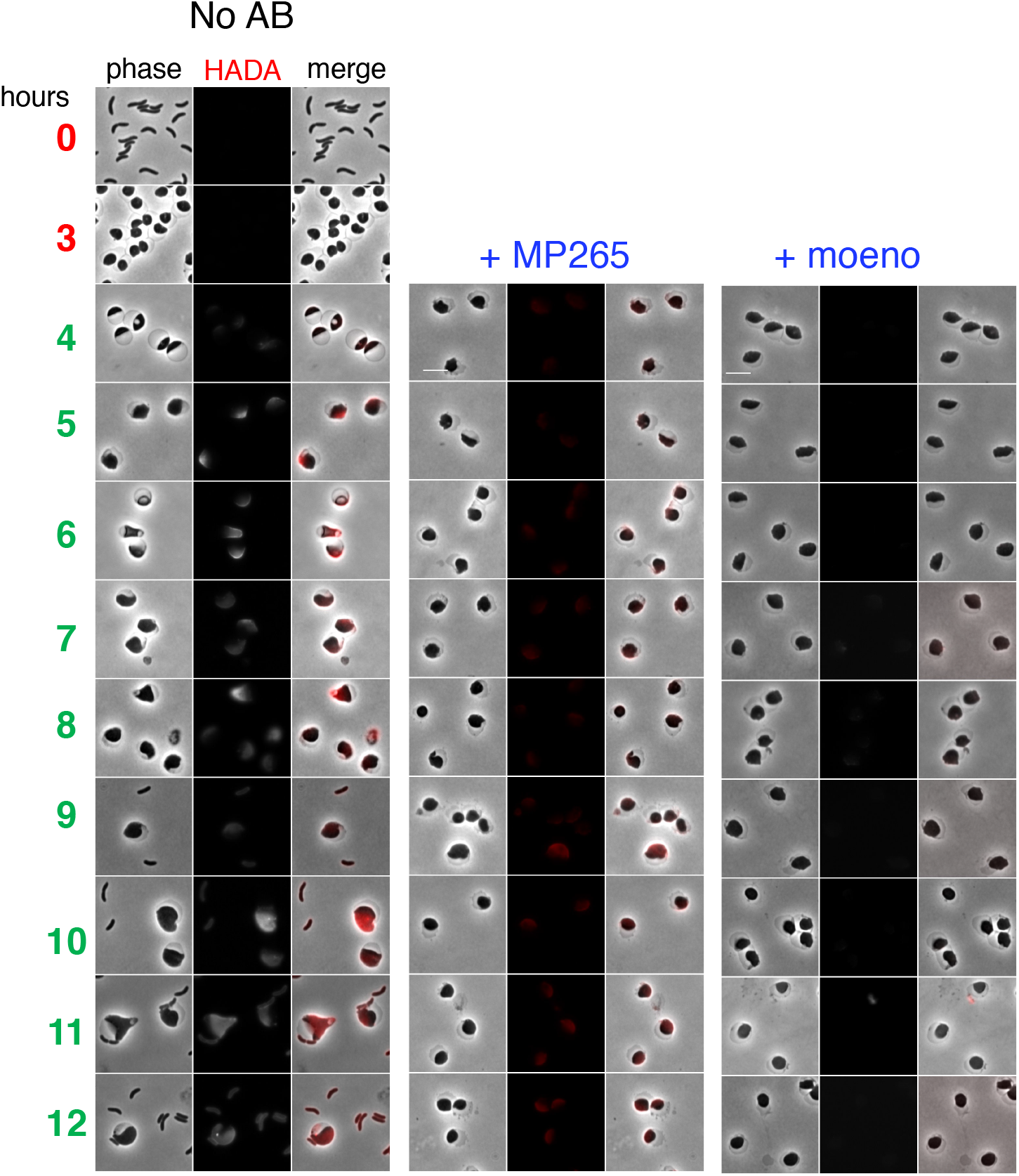
Deposition of new cell wall material in recovering spheres is primarily mediated by aPBPs. N16961 Δ*ldtA* Δ*ldtB* cells were exposed to PenG (100 μg ml^−1^, 10 × MIC) in M9 minimal medium for 3 h (T0 and T3), followed by washing and resuspension in fresh, pre-warmed M9 containing the fluorescent D-amino acid analog HADA as a cell wall label (4-12 h). Scale bar = 5 μm. The MreB inhibitor MP265 was added at 200 μM (10 × MIC) and the aPBP inhibitor moenomycin at 10 μg ml^−1^ (10x MIC).

### Identification of genes required for post-antibiotic recovery

We used transposon insertion site sequencing (TIS, aka TnSeq) to identify factors required for sphere recovery. Since we observed differences in recovery dynamics on solid (agarose pad) versus in liquid media, we combined both conditions in TIS experiments to uncover a broad array of recovery factors. Cells were exposed to PenG in liquid culture for 4 hr, washed, outgrown overnight in the absence of antibiotics, and then plated. The insertion sites in the Tn library were sequenced before addition of the antibiotic (PRE), after incubation in PenG (POST), and after the overnight outgrowth followed by plating (OG). Strikingly, comparison of the insertion profiles in the PRE and POST conditions (**Fig. 5A**) did not reveal any genes that met stringent criteria for differential fitness (>10-fold difference in insertion abundance, p-val <0.01). Thus, no single mutation appears to lead to catastrophic lysis in *V. cholerae* treated with penicillin G. In contrast, comparing the insertion profiles in the POST vs OG conditions revealed 55 genes which had reduced fitness during post-antibiotic outgrowth (>10-fold fewer insertions along with p-val <0.01).

**Figure 5.**
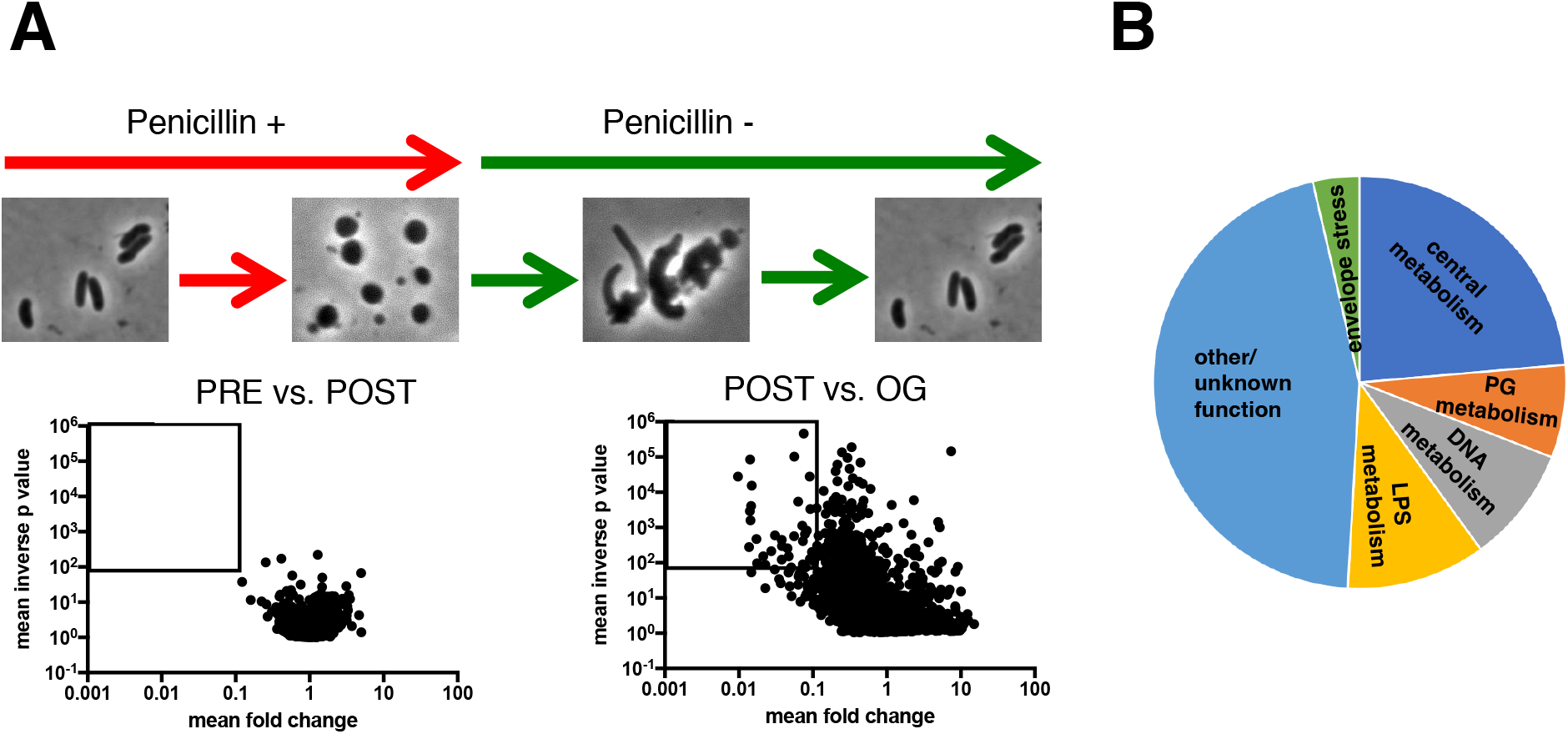
Identification of genes required for sphere recovery with TnSeq. **A**) Schematic of experimental design and volcano plots of change in relative abundance of insertion mutants between the two conditions (x axis) vs the concordance of independent insertion mutants within each gene (y axis, inverse p-value). The square denotes the cutoff criteria applied (>10-fold fitness defect, p-value <0.01) for identification of genes contributing to sphere recovery. PRE vs POST is a comparison of insertion frequencies after/before antibiotic exposure; POST vs. OG between an outgrowth period and directly after antibiotic exposure (see Methods for details) **B**) Distribution of the main functional categories of gene insertions that confer a post-antibiotic fitness defect. Functional categories were assigned manually following annotation of individual proteins in the Kegg database (http://www.genome.jp/kegg/).

Notably, there was an enrichment of several functional pathways among the 55 genes required for robust post-antibiotic outgrowth (**Fig. 5B**). Included among the enriched categories were genes predicted to be required for cell wall synthesis and recycling (*pbp1A, vc2153, ampG, pbp5, lpoB, mltA*), lipopolysaccharide (LPS) biosynthesis (core biosynthesis, Lipid A acylation and O-antigen synthesis pathways, *vc0212, vc0223, vc0225, vc0236, vc0237, vc0240*), intrinsic stress resistance (superoxide dismutase, *rpoE*), phosphate uptake (*vc0724-0726*) and chromosome dynamics (*mukBEF*) (**Fig. S3**).

Intriguingly, some of the hits (*vc2153, rpoE*, PG synthesis factors) were homologs or analogs of factors identified in a recent TnSeq screen for genes that promote tolerance to beta lactams in *Burkholderia pseudomallei* and *B. thailandensis* (6), raising the possibility that there are shared tolerance requirements across Gram-negative bacteria.

We were particularly interested in the contribution of cell envelope functions to sphere recovery and therefore prioritized genes involved in LPS and cell wall metabolism for further studies. We first focused on the 6 LPS biosynthesis genes that answered our screen. To validate the requirement of LPS core biosynthesis in the recovery process, we created an insertion mutant in *vc0225*, the gene encoding heptosyltransferase I. This mutation is expected to result in a truncated LPS molecule lacking an outer core and O-antigen and consistent with this, the mutant strain did not have detectable high molecular weight LPS in isolated outer membrane (OM) material (**Fig. S4**). Wild type (wt) and *vc0225::kan* mutant cells were compared in time-dependent viability experiments. Exposing wt cells to penicillin G (at 10 × MIC) in minimal medium (unlike LB) in some experiments permitted initial growth (**Fig. 6A**). We do not know the reason for this initial growth (*V. cholerae* does not become resistant to PenG in M9, as evidenced by sphere formation (**Fig. 6B**), but it is possible that antibiotic diffusion through the OM is slower in minimal than in rich medium. Disruption of the LPS core gene *vc0225* resulted in the absence of initial growth and a subsequent 100-fold plating defect after exposure to PenG (**Fig. 6A**), corroborating the TnSeq result. This survival defect could be complemented by expressing *vc0225* from a neutral chromosomal locus. Light microscopy revealed that the *vc0225* mutant strain still formed spheres (**Fig. 6B**); however, these spheres were morphologically distinct from wt spheres. In contrast to wt spheres, which were usually seen as single cells, exhibiting well-demarcated separation between the phase dark cytoplasm and the phase light periplasm (**Fig. 1A**), *vc0225* mutant cells were mostly grape-like masses showing a checkered pattern of distinct periplasmic enclaves in a sometimes divided cytoplasm (**Fig. 6B**). Visualizing an inner membrane marker (PBP1amCherry) also revealed the lack of a clear distinction between the inner and outer membrane in the mutant. Upon removal of the antibiotic, *vc0225* spheres were defective in all stages of the recovery process; the spheres underwent modest enlargement without initiating periplasmic engulfment (**Fig. 6C**). Thus, intact LPS appears necessary for sphere anatomy and internal organization; moreover, these LPS-associated sphere defects seem to impair sphere recovery.

**Figure 6.**
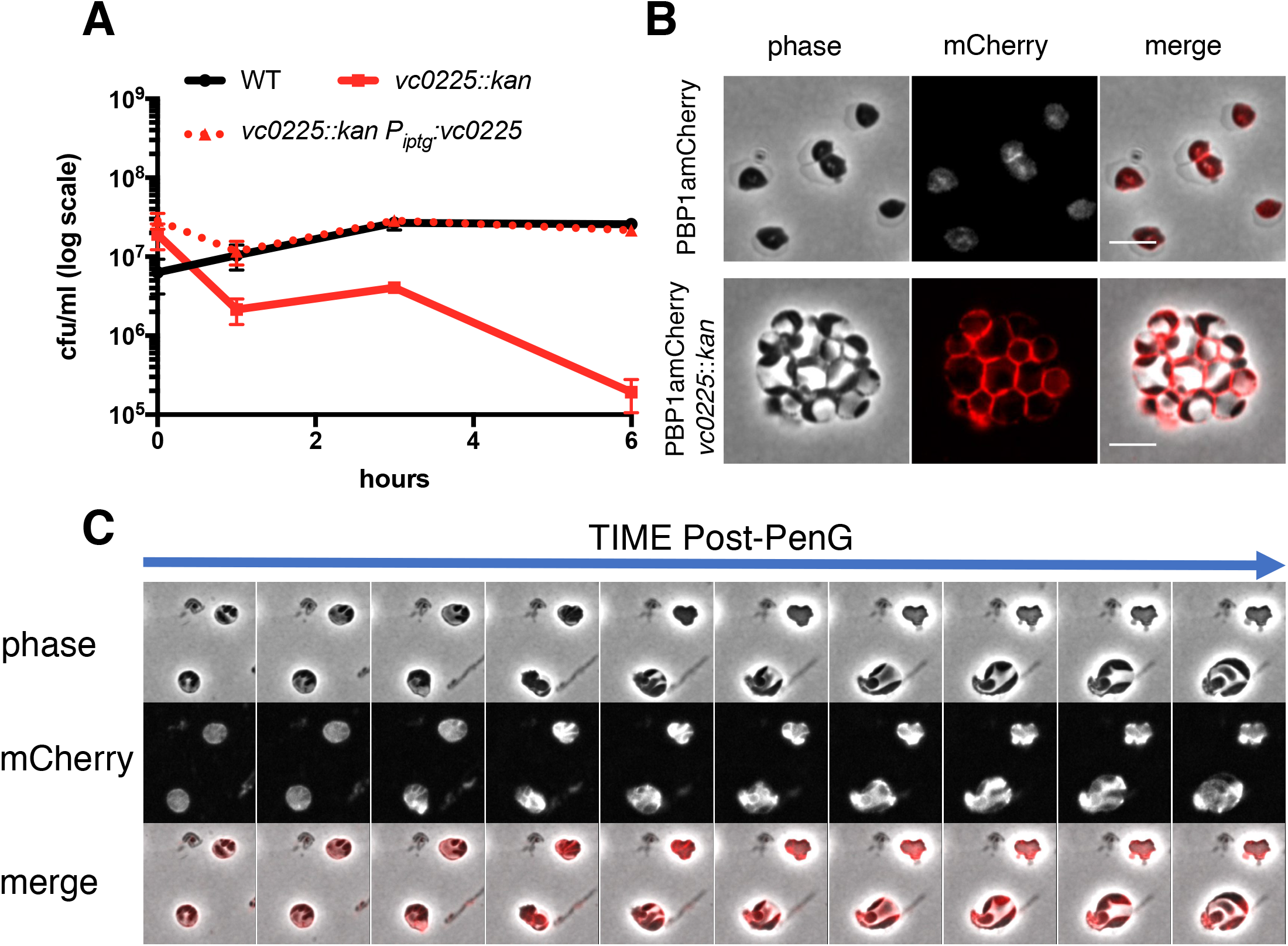
An LPS core mutant is defective in sphere recovery and organization. **A**) Time-dependent viability experiment in the presence of PenG. Strains were exposed to PenG (100 μg ml^−1^, 10x MIC) at T0 and plated for cfu ml^−1^ at the indicated times. **B**) Localization of Penicillin Binding Protein 1a (PBP1a) in PenG-treated wt (top) and mutant spheres (bottom). PBP1amCherry was expressed from its native, chromosomal locus. Scale bar = 5 μm **C**) Recovery pattern in a PenG-treated PBP1amCherry *vc0225* mutant after exposure to and subsequent removal of PenG. Frames are 10 min apart.

### Sphere integrity does not depend on Lipid A modifications

The outer membrane appears to be the principle load-bearing structure in beta lactam-induced spheres, because these cells are largely devoid of detectable cell wall material and are more susceptible to detergents and antimicrobial peptides (5, 7). *V. cholerae* LPS contains at least two modifications which are not found in *E. coli* LPS and that could potentially stabilize *V. cholerae* spheres. These modifications, addition of phosphoethanolamine to the 1-phosphate group of lipid A and an unusual glycine addition to a hydroxylauryl chain at the 2’ position of Lipid A, both promote resistance to polymyxin (26, 27); glycinylation (by the *alm* system) is dominant, but the pH-dependent EptA can promote residual polymyxin resistance when the *alm* system is inactivated (22). We investigated whether these modifications were required for sphere formation and integrity by deleting the *alm* operon, which encodes the glycine transferase activity, and *eptA*, which encodes the ethanolamine transferase. As expected, the *alm* mutation abrogated polymyxin resistance on LB (**Fig. S5**). When these mutants, either alone (not shown) or in combination, were exposed to PenG they formed spheres that were indistinguishable from wt spheres (**Fig. 7A**), indicating that these Lipid A modifications are not required for sphere generation. The Δalm Δ*eptA* mutations were then combined with disruptions in *vc0225* or *vc0212* (encoding the 3-hydroxy laurate transferase LpxN (28)) to test the effect of a core mutation (*vc0225*) or Lipid A under-acylation (*vc0212*) on sphere formation in LPS lacking glycine and ethanolamine modifications. (Note that mutation of *lpxN* results in the absence of the acyl chain modified by glycine and thus causes polymyxin B sensitivity, **Fig. S5**). The ΔalmΔeptA *vc0212::kan* mutant still formed spheres after PenG exposure (**Fig. 7A**) but these spheres had ~5-fold decrease in viability compared to the wt (**Fig. 7B**). The Δ*alm*Δ*eptA vc0225::kan* mutant also formed spheres, but resulted in a more dramatic reduction in viability compared to the Δ*alm*Δ*eptA vc0212::kan* mutant (**Fig. 7B**) and to the single *vc0225::kan* mutant, where there was less pronounced loss of viability after 3 h (compare with **Fig. 6A**). Thus, LPS core and O-antigen appear to be more critical for sphere formation/integrity and viability than Lipid A modifications. However, glycinylation and/or ethanolamine addition to Lipid A appear to promote maintenance of sphere integrity in the absence of LPS core and O-antigen, suggesting that these modifications contribute to OM stability in this context.

**Figure 7.**
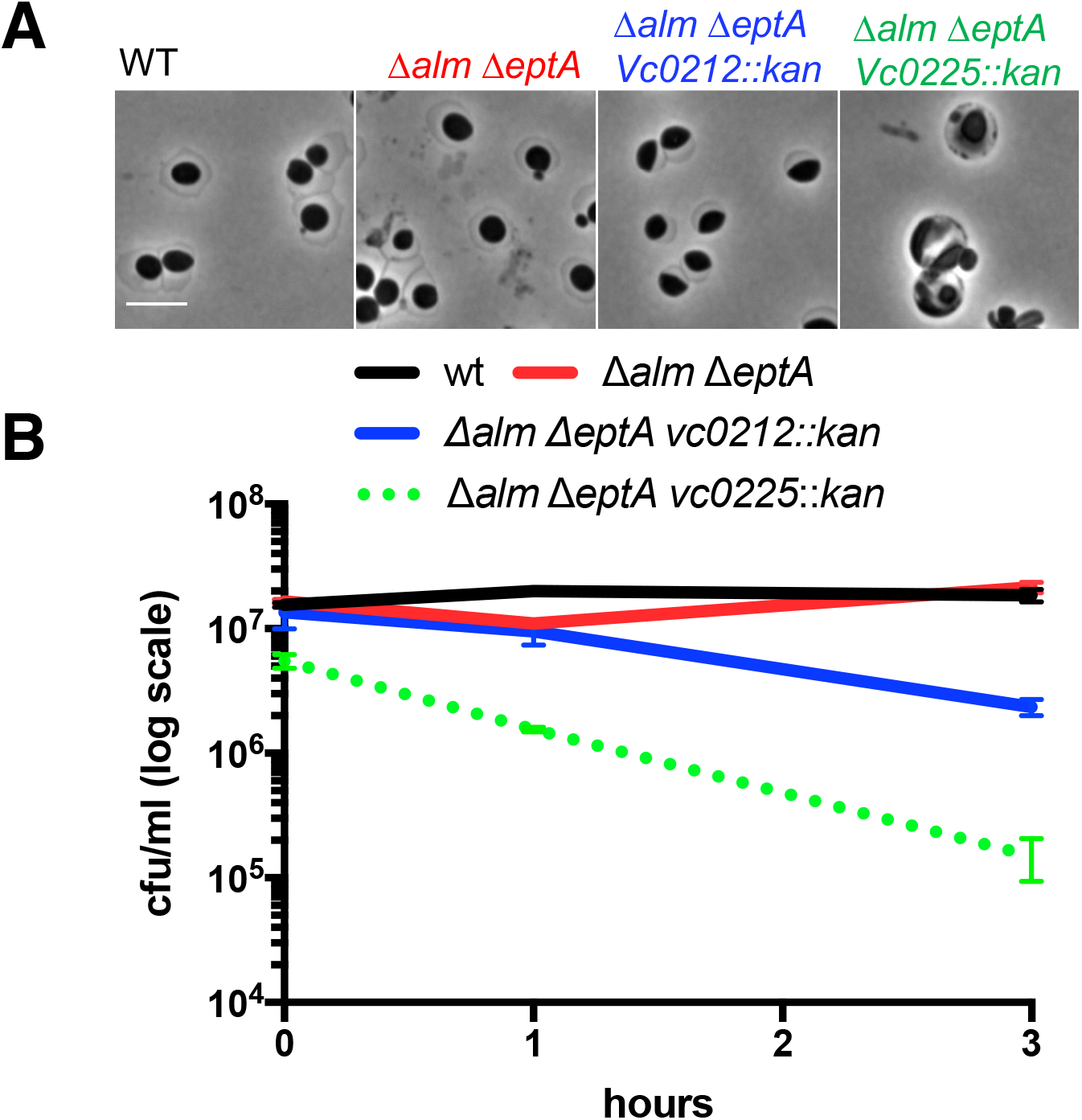
Outer membrane modifications are not required for sphere formation. **A**) Sphere formation in wt and mutants defective in glycine (*alm*) and ethanolamine (*eptA*) modifications alone and in combination with defects in LPS core biosynthesis (*vc0225*) and lipid A acylation (*vc0212*). Scale bar = 5 μm **B**) Time-dependent viability experiment as described in **Fig 4A**.

### The sigma E cell envelope stress response is required for penicillin tolerance

The TnSeq analysis implicated several genes in the sigma E cell envelope stress response as important for sphere viability/recovery (**Fig. S3**). Misfolding of outer membrane proteins such as OmpU triggers the *V. cholerae* envelope stress response, wherein sigma E directs the transcription of a set of genes involved in variety of cell envelope maintenance functions (29, 30). We found that the abundance of RpoE markedly increased several hours after cells were exposed to diverse antibiotics that interrupt cell wall synthesis (PenG, phosphomycin or D-cycloserine), consistent with the idea that this sigma factor promotes sphere survival (**Fig. 8A**). Interestingly, PenG treatment increased RpoE abundance independent of OmpU (**Fig. 8B**). The importance of sigma E for survival after PenG exposure was established by measuring time-dependent viability after antibiotic challenge. Since *rpoE* is essential, we used an Δ*rpoE*Δ*ompU* strain (the latter deletion enables *rpoE* deletion (30)), to investigate *rpoE’s* importance for sphere viability/recovery. Following exposure to PenG, the Δ*rpoE*Δ*ompU* strain exhibited a drastic (~1000-fold) plating defect compared to the wild type and Δ*ompU* controls (**Fig. 8B**). Thus, sigma E (and presumably the regulon it controls) is a crucial determinant of *V. cholerae* beta lactam tolerance.

**Figure 8.**
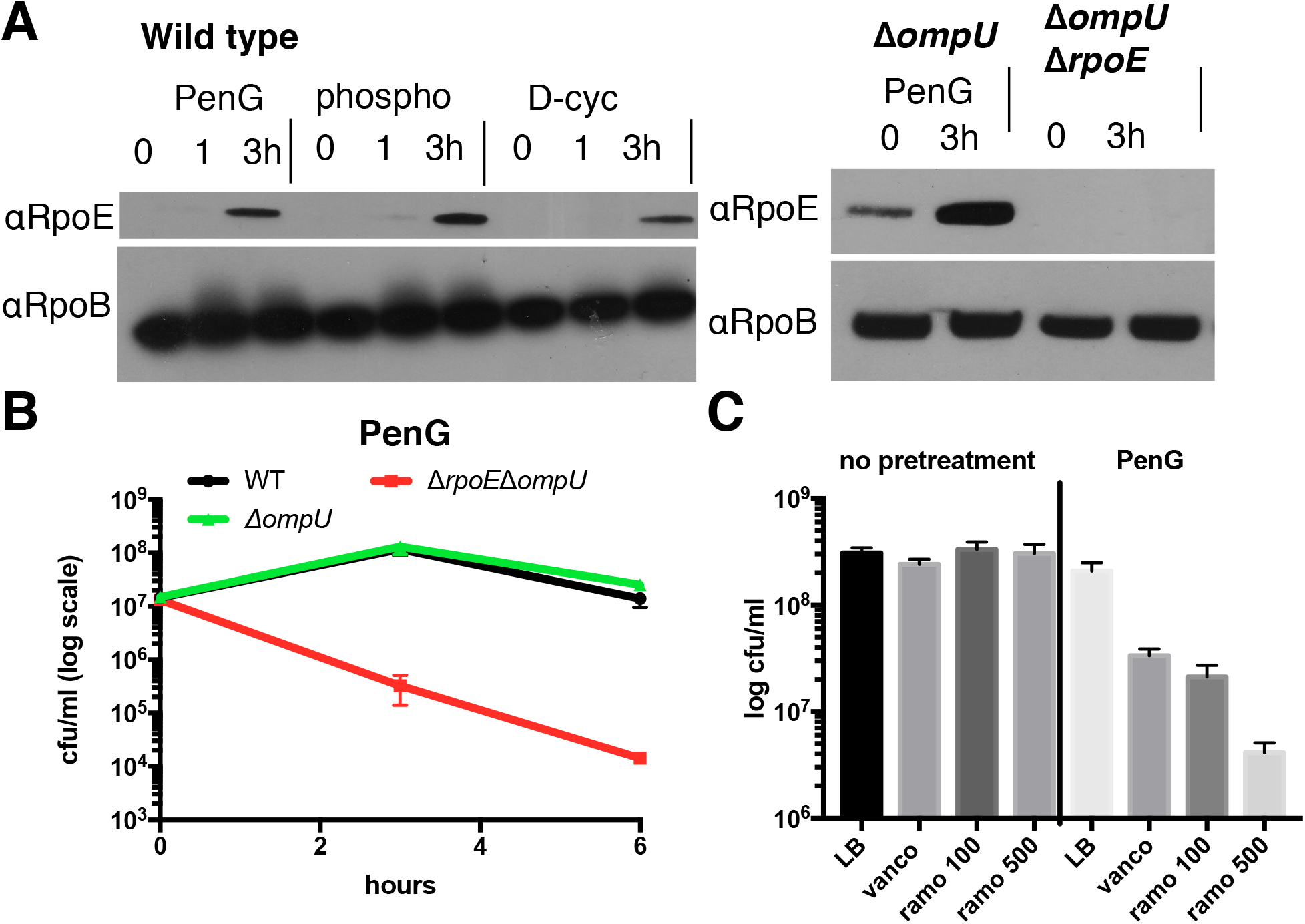
Sigma E is induced in response to cell wall acting antibiotics and is required for beta lactam tolerance. **A**) Western Blot using anti-RpoE antiserum after exposure to penicillin G (PenG, 100 μg ml^−1^, 10x MIC), phosphomycin (phospho, 100 μg ml^−1^, 3 × MIC) or D-cycloserine (D-cyc, 100 μg ml^−1^, 3 × MIC). **B**) Time-dependent viability of indicated strains after exposure to PenG (100 μg ml^−1^, 10 × MIC). **C**) Pretreatment with PenG sensitizes cells to high molecular weight antibiotics. Cells were exposed to either vehicle (no pretreatment) or Penicillin G (PenG, 100 μg ml^−1^) for 3h, followed by plating on either LB, vancomycin (vanco, 200 μg ml^−1^), or ramoplanin (ramo, 100 μg ml^−1^ or 500 μg ml^−1^).

The importance of the sigma E response for beta lactam tolerance indicated the strong possibility that PenG-treated *V. cholerae* sustain OM damage; this in turn suggested that beta lactam exposure might sensitize cells to high molecular weight (HMW) antibiotics that are typically too large to permeate the Gram-negative cell envelope; e.g. vancomycin and ramoplanin. To test this, we plated cells exposed to PenG or a vehicle control on either LB or plates containing vancomycin (100 μg ml^−1^) or ramoplanin (100 μg ml^−1^ and 500 μg ml^−1^) (**Fig. 8C**). While untreated cultures plated at close to 100% on any of these plates, pre-treatment with PenG for 3h resulted in a 10-to 50-fold plating defect on either HMW antibiotic. Thus, while *V. cholerae* is tolerant to beta lactam antibiotics, these agents appear to sensitize it against HMW antibiotics.

## Discussion

Antibiotic tolerance, the ability to survive and fully recover from exposure to normally lethal doses of bactericidal antibiotics, is a common cause of treatment failure and serves as a stepping-stone for the development of antibiotic resistance (31). The mechanism(s) of antibiotic tolerance and a related phenomenon, the post-antibiotic effect (the recovery process of tolerant cells) are understudied and insufficiently understood.

Here, we investigated the post-antibiotic effect in *V. cholerae*, an organism highly tolerant to typically bactericidal inhibitors of cell wall synthesis. Following beta lactam treatment, this pathogen forms viable cell wall deficient spheres that can re-establish their characteristic rod morphology when antibiotics are no longer present. We found that the process of restoration of rod shape appears to be media dependent and to differ from that described for the recovery of *E. coli* spheres induced by cefsulodin (19). At least on agarose pads, 3 successive steps characterized the recovery process. Recovery involves re-localization of PBP1a to the leading edge of a periplasmic engulfment process and aPBPs, rather than the Rod system, have a primary role, particularly in the early steps of recovery. The characteristic steps and protein localization patterns involved in restoration of rod shape suggests that *V. cholerae’s* ability to recover from a well-less spherical state may be a previously unappreciated type of programmed response to stress. Furthermore, the capacity of wall-less spheres to remain viable and to regain rod shape in a step-wise fashion is not restricted to *V. cholerae*, but found in several other Gram-negative organisms (7, 20, 32).

Not surprisingly, TnSeq analysis revealed that genes required for peptidoglycan biogenesis were essential for sphere recovery; however, additional envelope-related functions, particularly LPS biosynthesis, and the RpoE envelope stress response system were also critical for *V. cholerae* spheres to recover from beta lactam assault. While this TnSeq dataset offers many intriguing leads for future studies of the post-antibiotic recovery process (e.g. the role of riboflavin and condensins), here we primarily focused on the role of LPS and PG biosynthesis in role.

Mutations in LPS core and O-antigen biosynthesis can have at least two potentially detrimental consequences for sphere viability/recovery. First, truncated (rough) LPS has been shown to result in increased OM permeability due to the surface presentation of phospholipid bilayer patches resulting from LPS instability (33). Second, truncated LPS, which cannot be ligated to O-antigen, results in the accumulation of dead-end intermediates of O-antigen fused to the lipid carrier molecule undecaprenol (UNDP) (34). UNDP concentrations are limited and this carrier is critical for the biosynthesis of a variety of extracellular macromolecules, including peptidoglycan. Thus, in addition to direct effects on OM integrity, LPS core mutations can also impede efficient cell wall synthesis due to the reduced availability of UNDP; the consequences of such inhibition may be particularly severe for PG-deficient spheres. However, LPS core mutants exhibited post-antibiotic exposure phenotypes that cannot easily be explained by cell wall precursor depletion alone. The absence of the clear demarcation between the inner and outer membrane in the LPS core (*vc0225* mutant) deficient spheres suggests that IM material (IM phospholipids and associated proteins) may be present in the OM. More broadly, our findings demonstrate the importance of LPS integrity for *V. cholerae* survival of cell wall damage; it is likely that LPS structure/strength modulates susceptibility to beta lactams in other bacteria as well, as has been suggested in *E. coli* (35).

As expected, we also found cell wall synthesis and recycling factors to be required for sphere recovery. The prominent role of PBP1a, rather than its paralog PBP1b, in the recovery process supports our previous data showing that PBP1a is the principle aPBP in *V. cholerae* (23, 36) and is consistent with observations in lysozyme-treated, spherical *E. coli*, where that organism’s principal aPBP (PBP1B) is required for recovery (32). Intriguingly, PBP1a localized as a concentrated ring around the outgrowth area. Moenomycin, an aPBP TG inhibitor, abrogated PBP1a’s capacity to localize to the outgrowth area as well as sphere recovery, revealing the essentiality of aPBP enzymatic activity for the recovery process. In contrast, MreB, and by extension likely the associated Rod system appeared to play a minor role during the early stages of recovery (though MreB was ultimately necessary for full sphere-to-rod conversion). We do not know the exact structure of possible remnant PG in spheres; however, our results suggest that aPBPs are more efficient than the Rod system at starting PG synthesis in the absence of a rod-shaped scaffold. Recent data suggest that MreB determines the directionality of Rod-mediated PG synthesis through its axial, membrane-curvature-induced orientation in the cell (37). Our data are in line with such a model, as axial localization cues are lost in a sphere. Thus, the Rod system might rely on the aPBPs to first mediate some degree of sphere constriction, inducing heterogeneity in membrane curvature that would then enable ordered, Rod-mediated PG deposition resulting in cell elongation.

In summary, we provide here an analysis of factors required for post-antibiotic recovery in *V. cholerae* treated with a beta lactam antibiotic. Our results directly demonstrate a role for OM integrity in beta lactam tolerance and establish a differential role for aPBPs and the Rod system for post-antibiotic recovery. The factors identified here could serve as novel targets for antibiotic adjuvants that increase the efficacy of beta lactam antibiotics and other inhibitors of cell wall synthesis towards Gram-negative pathogens.

## Acknowledgements

We thank Hongbaek Cho and Thomas Bernhardt for the msfGFP template. Carol Gross is acknowledged for her gift of the anti RpoE antibody. Research in the Waldor lab is supported by the Howard Hughes Medical Institute and NIH (2R01-AI-042347-23). Research in the VanNieuwenhze lab is supported by NIH grant GM113172.

## Materials and Methods

### Media, chemicals and growth conditions

All growth experiments were conducted either in Luria Bertani Broth (LB) or M9 minimal medium supplemented with glucose (0.2 %). Antibiotics were purchased from the following suppliers: Moenomycin (Santa Cruz), MP265 (Santa Cruz) Penicillin G (Fisher), streptomycin (Santa Cruz). Unless otherwise noted, all experiments were conducted in three biological replicates (i.e. experiments conducted on different days). For each experiment, two independent overnight cultures were used; where possible (i.e. for mutants constructed in this study) these cultures were inoculated from two independently isolated clones.

### Molecular techniques/strain construction

For molecular cloning purposes, PCR was conducted using Q5 high fidelity polymerase (NEB). For diagnostic PCRs, OneTaq mastermix (NEB) was used instead. All cloned constructs were verified using sequencing. All plasmid cloning was done using Isothermal Assembly (ITA) (38). Oligos are summarized in **Table S1** and strains in **Table S2**. Unless otherwise noted, all strains were constructed in the El Tor N16961 background.

Unless otherwise noted, mutants were constructed as previously described using the suicide plasmid pCVD442 (39) and homologous recombination to replace genes with the sequence TAATGCGGCCGCACTCGAGTAATAATGATGA. Briefly, the *E. coli* donor strain Sm10 carrying a pCVD-based deletion plasmid was mixed 1:1 with the *V. cholerae* recipient on an LB plate and incubated for at least 2h at 37 °C. The cell mixture was then streaked out on a plate containing carbenicillin (100 μg/ml) and streptomycin (200 μg/ml) to select against the donor strain and for recipients that have integrated the deletion plasmid. To counterselect against pCVD, a single colony was then streaked out on sucrose agar (15 g/L agar, 10 g/L tryptone, 5 g/L yeast extract, filter-sterilized sucrose added after autoclaving to 10 % final concentration) + streptomycin. Plates were incubated at ambient temperature for 1 day and then transferred to 30 °C, followed by additional incubation for 1 or 2 days. Successful mutants were verified via colony PCR using primers flanking the gene of interest.

For the *vc0225* disruption, sacB-based counterselection did not work due to the inability of LPS core mutants to grow on sucrose agar. We therefore used a kanR (conferring kanamycin reistance) variant of the single crossover suicide vector pGP704 (40) to create insertion disruption mutants. To this end, a 400bp internal fragment (nucleotide position 27 – 427) of *vc0225* or a 461 bp internal fragment (nucleotide position 93 – 554) of *vc0212* was cloned into pGPkan and transferred into recipient *V. cholerae* using the di-amino pimelic acid (DAP)-auxotrophic *E. coli* donor strain MFD lambda pir (41). Insertion mutants were selected on plates containing streptomycin (200 μg/ml) and kanamycin (50 μg/ml).

Procedures for the construction of other knockout plasmids and strains was as follows:

#### Pbp1a::pbp1amcherry

Upstream and downstream regions were amplified using primers TDP 1362/1434 and TDP 1435/232 respectively and fused with mCherry (amplified using primers 1436/1437) and Xba1-digested pCVD442 using ITA. These primers insert the linker sequence gacatcctcgagctc between PBP1a (no stop codon) and mCherry. Δ*alm* Δ*eptA*

For the Δ*alm* plasmid, upstream and downstream homologies were amplified using primers 519/520 and 521/522 respectively. For Δ*eptA*, upstream and downstream homology regions were amplified using primers 539/540 and 541/542.

### TnSeq

TnSeq was conducted as described before (42–44); briefly, cultures were subjected to transposon (mariner) mutagenesis in duplicate. In the resulting libraries (~200,000 colonies/replicate), whle population transposon-chromosome junctions were sequenced (PRE sample, see below); the libraries were then frozen down in 30 % glycerol (−80°C). For the experiment, libraries were grown to an OD_600_ ~0.5 then exposed to penicillin G (100 μg/ml, 10 × MIC) for 4 h and sequenced again (POST sample), followed by washing to remove the antibiotic and outgrowth overnight; after which the libraries were sequenced again (outgrowth, OG sample). Sequencing was performed as follows. Pelleted libraries were lysed and DNA fragmented using NEB fragmentase, followed by blunting (Blunting enzyme mix, NEB), A-tailing and ligation of specific adaptors. Transposon-DNA junctions were then PCR amplified using transposon- and adaptor-specific primers. The libraries were then sequenced on an Illumina MiSeq. Data analysis was essentially conducted as described (42–44), however, to avoid false negatives that did not pass our stringent cut-off, we also used a candidate-based approach (based on known genetic interactions between cell envelope functions) to visually inspect the TnSeq dataset in the genome browser Artemis (this approach yielded e.g. WigKR and RpoE).

### Recovery on agarose pads

For recovery timelapses, overnight cultures were diluted 100-fold into fresh M9 MM, then grown until OD600 = 0.3 (3.5 h) at 37 °C shaking. Antibiotic was then added, followed by incubation for another 3h. Cells were then washed twice in antibiotic-free medium and applied to agarose patches (0.8 % agarose in M9) and imaged every 5 min on a Leica DMi8 inverted microscope with incubated (30 °C) stage. For fluorescent readings, exposure time was 500 ms (mCherry), 300 ms (msfGFP) or 1000 ms (HADA).

### Recovery in liquid medium

Overnight cultures were diluted 100fold into fresh M9 MM, then grown until OD600 = 0.3 (3.5 h) at 37 °C shaking. Antibiotic was then added, followed by incubation for another 3h. Cells were then washed twice in antibiotic-free medium and diluted 10fold into same containing 100 μM HADA. Cells were withdrawn at the indicated time points, washed once with M9 and imaged as detailed above.

---

**Figure S1.**
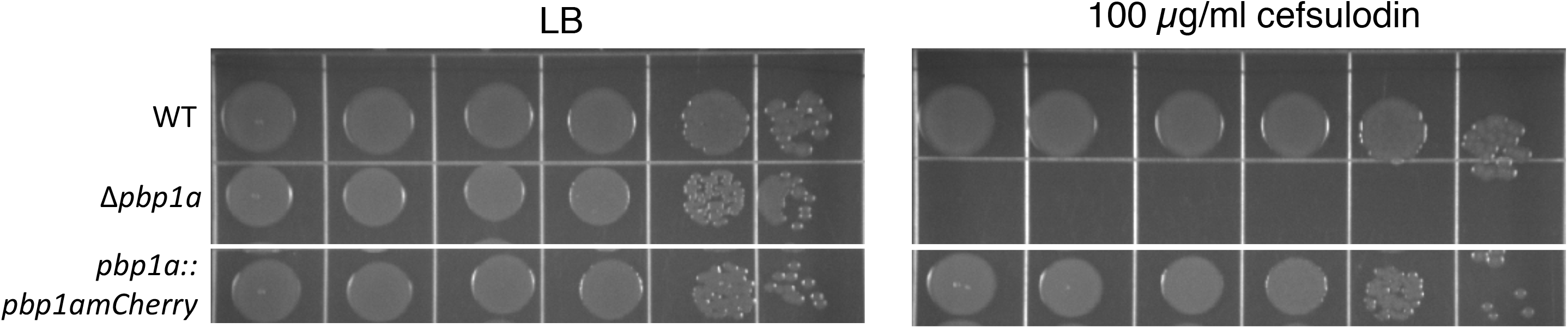
Fluorescent protein fusion to PBP1a is functional. Overnight cultures of the indicated strains were plated on LB agar containing 100 μg ml^−1^ cefsulodin, an inhibitor of PBP1B (23).

**Figure S2.**
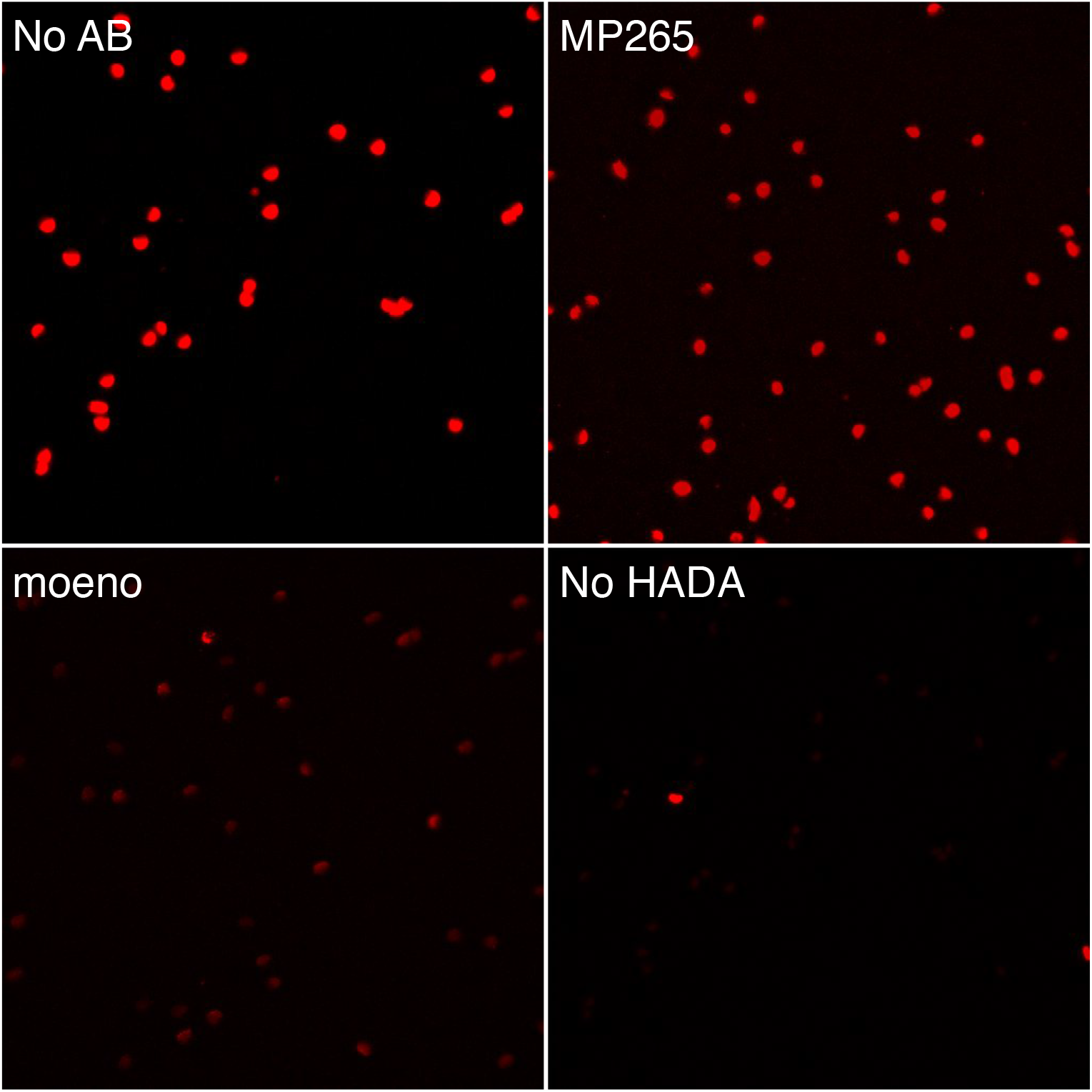
HADA staining in recovering spheres treated with inhibitors of aPBPs or the Rod system. Frames are taken from the experiment as shown in **Figure 4**. The pixel intensity range was set to 100 – 2000 in all images to illustrate the above background fluorescence in moenomycin-treated cells. Note that this results in overexposure of the untreated sample image.

**Figure S3.**
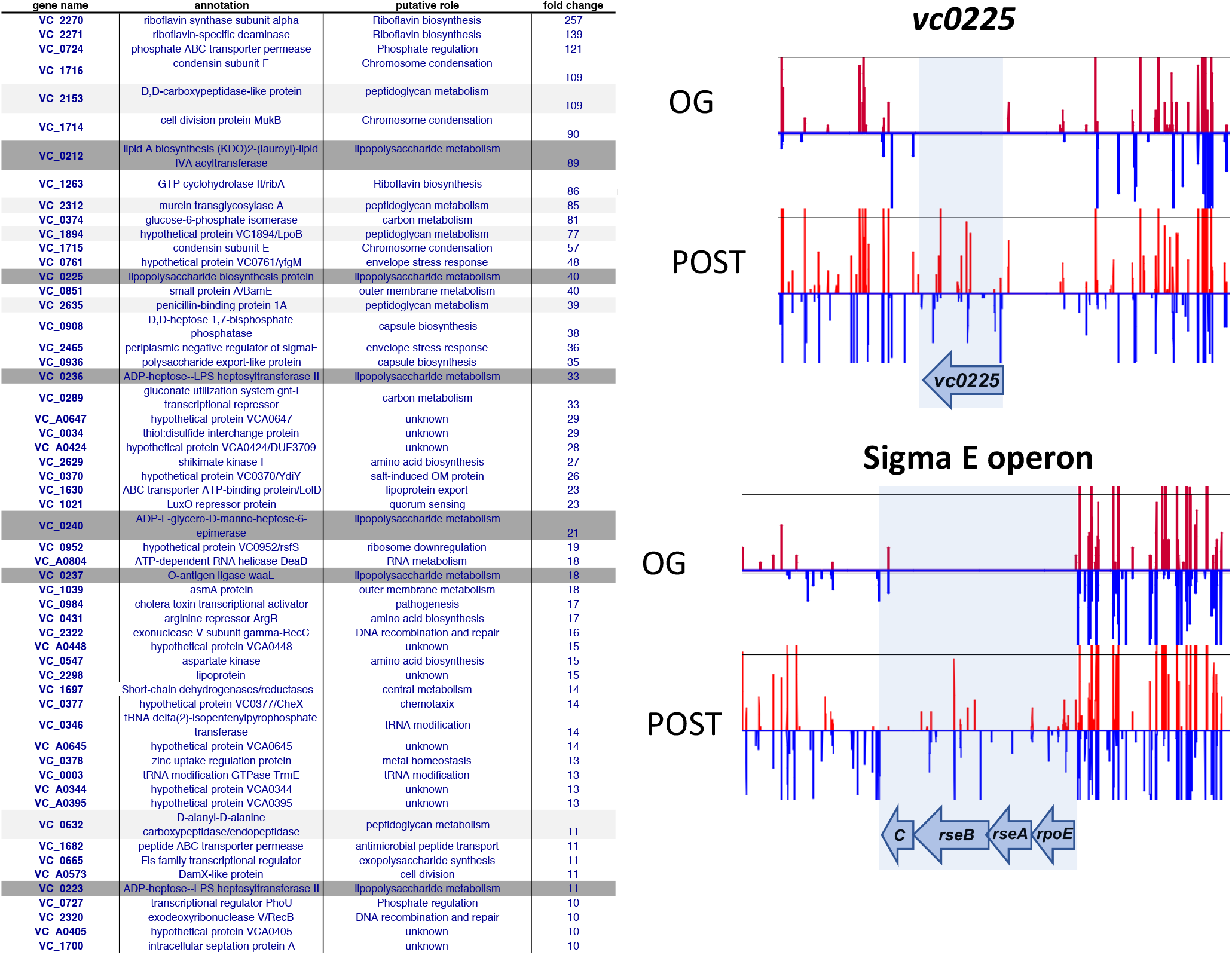
**A**) TnSeq hits below the cutoff (>/= 10-fold fitness defect, p-val < 0.01). **B**) representative Artemis plots of the genes encoding Heptosyltransferase I (*vc0225*) and the Sigma E operon. POST results from analyses Post-antibiotic; OG results from analyses after post-antibiotic outgrowth.

**Figure S4.**
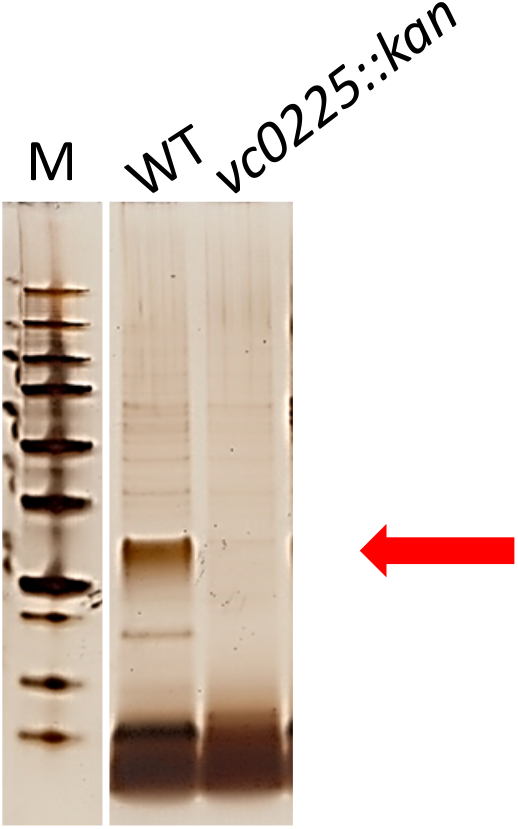
Silver stain of *V. cholerae* isolated outer membranes, comparing wild type and the *vc0225* mutant. The red arrow points to high molecular weight structures, likely core + O-antigen, M, marker.

**Figure S5.**
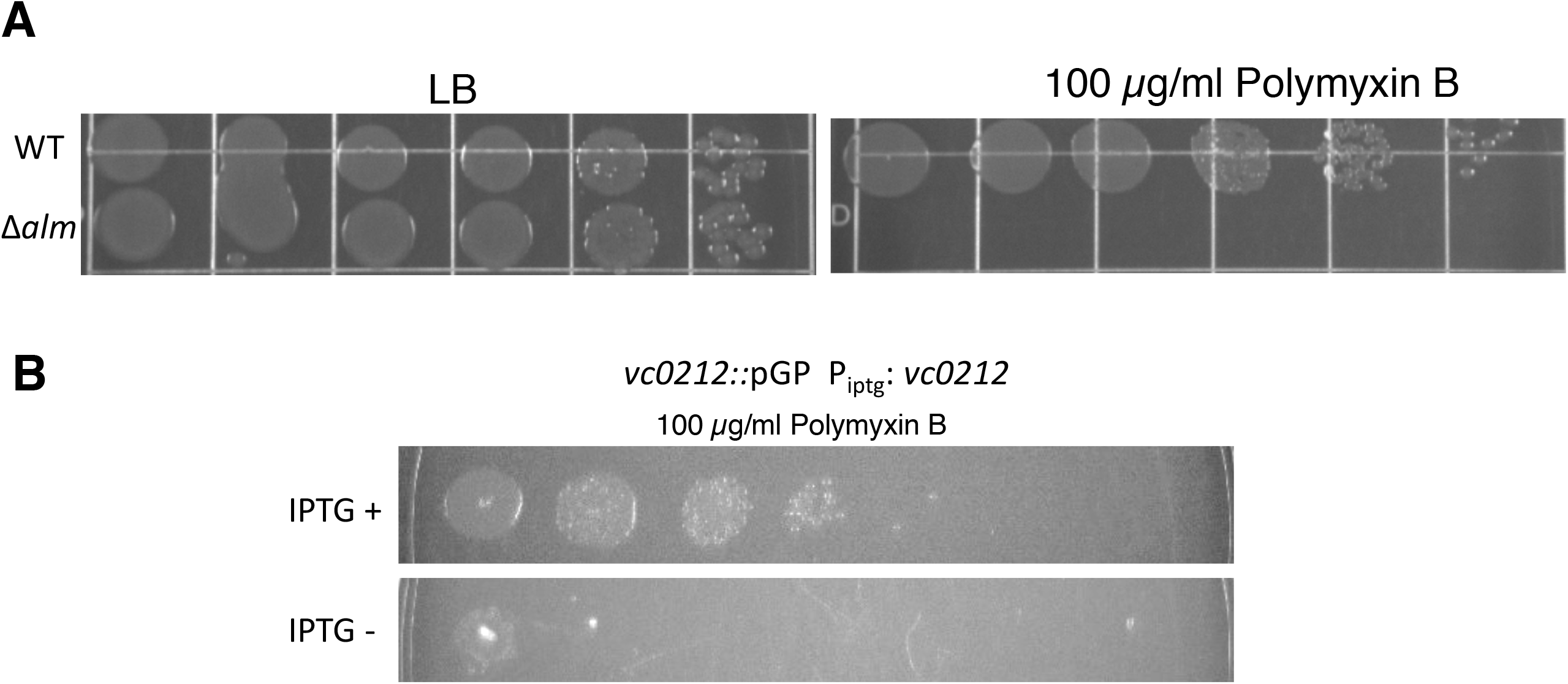
Polymyxin sensitivity of *alm* and *eptA* mutants. Overnight cultures of the indicated strains were plated on LB agar containing Polymyxin B at 10 μg ml^−1^ and IPTG (100 μM) where applicable.

**Movie S1.** Representative timelapse of wt *V. cholerae* sphere recovery. Cells were exposed to PenG in M9 MM + glucose, followed by washing and transfer to an agarose pad containing M9 MM + glucose. Frames are 5 min apart.

## References

1. Meylan S, Andrews IW, Collins JJ. 2018. Targeting Antibiotic Tolerance, Pathogen by Pathogen. Cell 172:1228–1238.

2. Conlon BP, Rowe SE, Lewis K. 2015. Persister cells in biofilm associated infections. Adv Exp Med Biol 831:1–9.

3. Sharma B, Brown AV, Matluck NE, Hu LT, Lewis K. 2015. Borrelia burgdorferi, the Causative Agent of Lyme Disease, Forms Drug-Tolerant Persister Cells. Antimicrob Agents Chemother 59:4616–24.

4. Lewis K. 2010. Persister cells. Annu Rev Microbiol 64:357–72.

5. Dorr T, Davis BM, Waldor MK. 2015. Endopeptidase-mediated beta lactam tolerance. PLoS Pathog 11:e1004850.

6. Held K, Gasper J, Morgan S, Siehnel R, Singh P, Manoil C. 2018. Determinants of Extreme beta-Lactam Tolerance in the Burkholderia pseudomallei Complex. Antimicrob Agents Chemother 62.

7. Monahan LG, Turnbull L, Osvath SR, Birch D, Charles IG, Whitchurch CB. 2014. Rapid conversion of Pseudomonas aeruginosa to a spherical cell morphotype facilitates tolerance to carbapenems and penicillins but increases susceptibility to antimicrobial peptides. Antimicrob Agents Chemother 58:1956–62.

8. Errington J, Mickiewicz K, Kawai Y, Wu LJ. 2016. L-form bacteria, chronic diseases and the origins of life. Philos Trans R Soc Lond B Biol Sci 371.

9. Dorr T, Alvarez L, Delgado F, Davis BM, Cava F, Waldor MK. 2016. A cell wall damage response mediated by a sensor kinase/response regulator pair enables beta-lactam tolerance. Proc Natl Acad Sci U S A 113:404–9.

10. Cheng AT, Ottemann KM, Yildiz FH. 2015. Vibrio cholerae Response Regulator VxrB Controls Colonization and Regulates the Type VI Secretion System. PLoS Pathog 11:e1004933.

11. Teschler JK, Cheng AT, Yildiz FH. 2017. The Two-Component Signal Transduction System VxrAB Positively Regulates Vibrio cholerae Biofilm Formation. J Bacteriol 199.

12. Zhao H, Patel V, Helmann JD, Dorr T. 2017. Don’t let sleeping dogmas lie: new views of peptidoglycan synthesis and its regulation. Mol Microbiol 106:847–860.

13. Typas A, Banzhaf M, van den Berg van Saparoea B, Verheul J, Biboy J, Nichols RJ, Zietek M, Beilharz K, Kannenberg K, von Rechenberg M, Breukink E, den Blaauwen T, Gross CA, Vollmer W. 2010. Regulation of peptidoglycan synthesis by outer-membrane proteins. Cell 143:1097–109.

14. Paradis-Bleau C, Markovski M, Uehara T, Lupoli TJ, Walker S, Kahne DE, Bernhardt TG. 2010. Lipoprotein cofactors located in the outer membrane activate bacterial cell wall polymerases. Cell 143:1110–20.

15. Cho H, Wivagg CN, Kapoor M, Barry Z, Rohs PD, Suh H, Marto JA, Garner EC, Bernhardt TG. 2016. Bacterial cell wall biogenesis is mediated by SEDS and PBP polymerase families functioning semi-autonomously. Nat Microbiol doi:10.1038/nmicrobiol.2016.172:16172.

16. Garner EC, Bernard R, Wang W, Zhuang X, Rudner DZ, Mitchison T. 2011. Coupled, circumferential motions of the cell wall synthesis machinery and MreB filaments in B. subtilis. Science 333:222–5.

17. Dominguez-Escobar J, Chastanet A, Crevenna AH, Fromion V, Wedlich-Soldner R, Carballido-Lopez R. 2011. Processive movement of MreB-associated cell wall biosynthetic complexes in bacteria. Science 333:225–8.

18. van Teeffelen S, Wang S, Furchtgott L, Huang KC, Wingreen NS, Shaevitz JW, Gitai Z. 2011. The bacterial actin MreB rotates, and rotation depends on cell-wall assembly. Proc Natl Acad Sci U S A 108:15822–7.

19. Billings G, Ouzounov N, Ursell T, Desmarais SM, Shaevitz J, Gitai Z, Huang KC. 2014. De novo morphogenesis in L-forms via geometric control of cell growth. Mol Microbiol 93:883–96.

20. Ranjit DK, Young KD. 2013. The Rcs stress response and accessory envelope proteins are required for de novo generation of cell shape in Escherichia coli. J Bacteriol 195:2452–62.

21. Morgenstein RM, Bratton BP, Nguyen JP, Ouzounov N, Shaevitz JW, Gitai Z. 2015. RodZ links MreB to cell wall synthesis to mediate MreB rotation and robust morphogenesis. Proc Natl Acad Sci U S A 112:12510–5.

22. Lee TK, Meng K, Shi H, Huang KC. 2016. Single-molecule imaging reveals modulation of cell wall synthesis dynamics in live bacterial cells. Nat Commun 7:13170.

23. Dorr T, Moll A, Chao MC, Cava F, Lam H, Davis BM, Waldor MK. 2014. Differential requirement for PBP1a and PBP1b in in vivo and in vitro fitness of Vibrio cholerae. Infect Immun 82:2115–24.

24. Takacs CN, Poggio S, Charbon G, Pucheault M, Vollmer W, Jacobs-Wagner C. 2010. MreB drives de novo rod morphogenesis in Caulobacter crescentus via remodeling of the cell wall. J Bacteriol 192:1671–84.

25. Cava F, de Pedro MA, Lam H, Davis BM, Waldor MK. 2011. Distinct pathways for modification of the bacterial cell wall by non-canonical D-amino acids. EMBO J 30:3442–53.

26. Herrera CM, Henderson JC, Crofts AA, Trent MS. 2017. Novel coordination of lipopolysaccharide modifications in Vibrio cholerae promotes CAMP resistance. Mol Microbiol 106:582–596.

27. Hankins JV, Madsen JA, Giles DK, Brodbelt JS, Trent MS. 2012. Amino acid addition to Vibrio cholerae LPS establishes a link between surface remodeling in gram-positive and gram-negative bacteria. Proc Natl Acad Sci U S A 109:8722–7.

28. Hankins JV, Madsen JA, Giles DK, Childers BM, Klose KE, Brodbelt JS, Trent MS. 2011. Elucidation of a novel Vibrio cholerae lipid A secondary hydroxy-acyltransferase and its role in innate immune recognition. Mol Microbiol 81:1313–29.

29. Mathur J, Davis BM, Waldor MK. 2007. Antimicrobial peptides activate the Vibrio cholerae sigmaE regulon through an OmpU-dependent signalling pathway. Mol Microbiol 63:848–58.

30. Davis BM, Waldor MK. 2009. High-throughput sequencing reveals suppressors of Vibrio cholerae rpoE mutations: one fewer porin is enough. Nucleic Acids Res 37:5757–67.

31. Levin-Reisman I, Ronin I, Gefen O, Braniss I, Shoresh N, Balaban NQ. 2017. Antibiotic tolerance facilitates the evolution of resistance. Science 355:826–830.

32. Ranjit DK, Jorgenson MA, Young KD. 2017. PBP1B Glycosyltransferase and Transpeptidase Activities Play Different Essential Roles during the De Novo Regeneration of Rod Morphology in Escherichia coli. J Bacteriol 199.

33. Malinverni JC, Silhavy TJ. 2009. An ABC transport system that maintains lipid asymmetry in the gram-negative outer membrane. Proc Natl Acad Sci U S A 106:8009–14.

34. Jorgenson MA, Young KD. 2016. Interrupting Biosynthesis of O Antigen or the Lipopolysaccharide Core Produces Morphological Defects in Escherichia coli by Sequestering Undecaprenyl Phosphate. J Bacteriol 198:3070–3079.

35. Yao Z, Kahne D, Kishony R. 2012. Distinct single-cell morphological dynamics under beta-lactam antibiotics. Mol Cell 48:705–12.

36. Dorr T, Lam H, Alvarez L, Cava F, Davis BM, Waldor MK. 2014. A novel peptidoglycan binding protein crucial for PBP1A-mediated cell wall biogenesis in Vibrio cholerae. PLoS Genet 10:e1004433.

37. Hussain S, Wivagg CN, Szwedziak P, Wong F, Schaefer K, Izore T, Renner LD, Holmes MJ, Sun Y, Bisson-Filho AW, Walker S, Amir A, Lowe J, Garner EC. 2018. MreB filaments align along greatest principal membrane curvature to orient cell wall synthesis. Elife 7.

38. Gibson DG, Young L, Chuang RY, Venter JC, Hutchison CA, 3rd, Smith HO. 2009. Enzymatic assembly of DNA molecules up to several hundred kilobases. Nat Methods 6:343–5.

39. Donnenberg MS, Kaper JB. 1991. Construction of an eae deletion mutant of enteropathogenic Escherichia coli by using a positive-selection suicide vector. Infect Immun 59:4310–7.

40. Miller VL, Mekalanos JJ. 1988. A novel suicide vector and its use in construction of insertion mutations: osmoregulation of outer membrane proteins and virulence determinants in Vibrio cholerae requires toxR. J Bacteriol 170:2575–83.

41. Ferrieres L, Hemery G, Nham T, Guerout AM, Mazel D, Beloin C, Ghigo JM. 2010. Silent mischief: bacteriophage Mu insertions contaminate products of Escherichia coli random mutagenesis performed using suicidal transposon delivery plasmids mobilized by broad-host-range RP4 conjugative machinery. J Bacteriol 192:6418–27.

42. Yamaichi Y, Dorr T. 2017. Transposon Insertion Site Sequencing for Synthetic Lethal Screening. Methods Mol Biol 1624:39–49.

43. Pritchard JR, Chao MC, Abel S, Davis BM, Baranowski C, Zhang YJ, Rubin EJ, Waldor MK. 2014. ARTIST: high-resolution genome-wide assessment of fitness using transposon-insertion sequencing. PLoS Genet 10:e1004782.

44. Chao MC, Abel S, Davis BM, Waldor MK. 2016. The design and analysis of transposon insertion sequencing experiments. Nat Rev Microbiol 14:119–28.

